# Airway IRF7^hi^ versus IRF7^lo^ molecular response patterns determine clinical phenotypes in children with acute wheezing

**DOI:** 10.1101/222950

**Authors:** Siew-Kim Khoo, James Read, Kimberley Franks, Guicheng Zhang, Joelene Bizzintino, Laura Coleman, Christopher McCrae, Lisa Öberg, Niamh Troy, Franciska Prastanti, Janet Everard, Stephen Oo, Meredith L Borland, Rose A Maciewicz, Peter N Le Souëf, Ingrid A Laing, Anthony Bosco

## Abstract

Asthma exacerbations are triggered by rhinovirus infections. We employed a systems biology approach to delineate upper airway gene network patterns underlying asthma exacerbation phenotypes in children. Cluster analysis unveiled distinct IRF7^hi^ versus IRF7^lo^ molecular phenotypes, the former exhibiting robust upregulation of Th1/type I interferon responses and the latter an alternative signature marked by upregulation of cytokine and growth factor signalling and downregulation of interferon gamma. The two phenotypes also produced distinct clinical phenotypes. For IRF7^lo^ versus IRF7^hi^: symptom duration prior to hospital presentation was more than twice as long from initial symptoms (p=0.011) and nearly three times as long for cough (p<0.001); the odds ratio of admission to hospital was increased more than four-fold (p=0.018); and time to recurrence was shorter (p=0.015). In summary, our findings demonstrate that asthma exacerbations in children can be divided into IRF7^hi^ versus IRF7^lo^ phenotypes with associated differences in clinical phenotypes.

**Abbreviations:** AHR, airway hyperresponsiveness; ARG1, Arginase 1, CSF3, Colony Stimulating Factor 3; CD38, Cluster of Differentiation 38; CD163, Cluster of Differentiation 163; cDCs, conventional (or myeloid) dendritic cells; DDX60, DExD/H-Box Helicase 60; ED, Emergency Department; EGF, Epidermal Growth Factor; ERK, Extracellular signal-Regulated Kinase; FCER1G, Fc Fragment Of IgE Receptor Ig; HMBS, Hydroxymethylbilane Synthase; IFNg, Interferon Gamma; IFNL1, Interferon Lambda 1; IL-1R2, Interleukin 1 Receptor Type 2; IRF7, Interferon Regulatory Factor 7; ISG15, Interferon-stimulated gene 15; MDA5, Melanoma Differentiation-Associated protein 5; MX1, Myxovirus Resistance Protein 1; NAD, nicotinamide adenine dinucleotide; NCR1, Natural cytotoxicity triggering receptor 1; OSM, Oncostatin M; PD-L1, Programmed Death-Ligand 1; PPIA, Peptidylprolyl Isomerase A; PPIB Peptidylprolyl Isomerase B; RSAD2, Radical S-adenosyl methionine domain-containing protein 2; RSV, respiratory syncytial virus; RT-qPCR, quantitative reverse transcription PCR; RV, rhinovirus; sPLA2, secretory Phospholipase A2; TGFb, Transforming Growth Factor beta; THBS1, Thrombospondin 1; TNF, Tumor Necrosis Factor; TLR2, Toll-like Receptor 2.

## Introduction

Exacerbations of asthma and wheeze are mostly triggered by respiratory viral infections and are one of the most common reasons for a child to seek emergency care.^1^ Previous studies from our group have demonstrated that rhinovirus (RV) species C is the most frequent viral pathogen detected in children who present to the local emergency department (ED) with an asthma exacerbation.^2^ However, the molecular mechanisms that determine susceptibility to RV and expression of respiratory symptoms are not well understood. Previous investigations of airway epithelial cells infected with RV *in vitro* found that type I and III interferon responses were deficient in adults with asthma, leading to impaired viral control and exaggerated secondary responses.^3, 4^ However, this finding was replicated in some studies but not others.^5^ Human adult asthmatic volunteers experimentally infected with RV *in vivo* have exaggerated IL-25 and IL-33 responses, which drive Th2 inflammation.^6, 7^ Although these data provide a plausible mechanism to link RV infection with the pathogenesis of asthma, they are based on experimental models with artificial infections from laboratory RV strains. Hence, the extent they can recreate the complex environmental conditions underpinning natural RV-induced exacerbations is unclear.^1, 8^ Indeed, studies of naturally occurring virus-induced exacerbations report increased rather than decreased interferon responses.^9–11^ In this study, we have utilized an unbiased, systems biology approach to elucidate the innate immune mechanisms that are operating in the upper airways during natural exacerbations of asthma or wheezing in children.^9, 10^ Our findings provide new insights into the role of gene networks, particularly IRF7, and their relationship to clinical phenotypes in this disease.

## Results

### Characteristics of the study population

The study was based on a case/control design. The cases (n=56) consisted of children who presented to the ED with an acute exacerbation of asthma or wheeze. The controls consisted of children who were either siblings of the cases, or they were recruited from the general community (controls, n=31). Convalescent samples (n=19) were available from children after they had recovered from an acute exacerbation of asthma or wheeze, but only a subset of these convalescent samples (n=5/19) were paired with acute samples. Samples from an independent group of children (n=99) with exacerbations of asthma or wheeze were utilised as a replication cohort. The characteristics of the study participants are presented in Table 1.

**Table 1:**
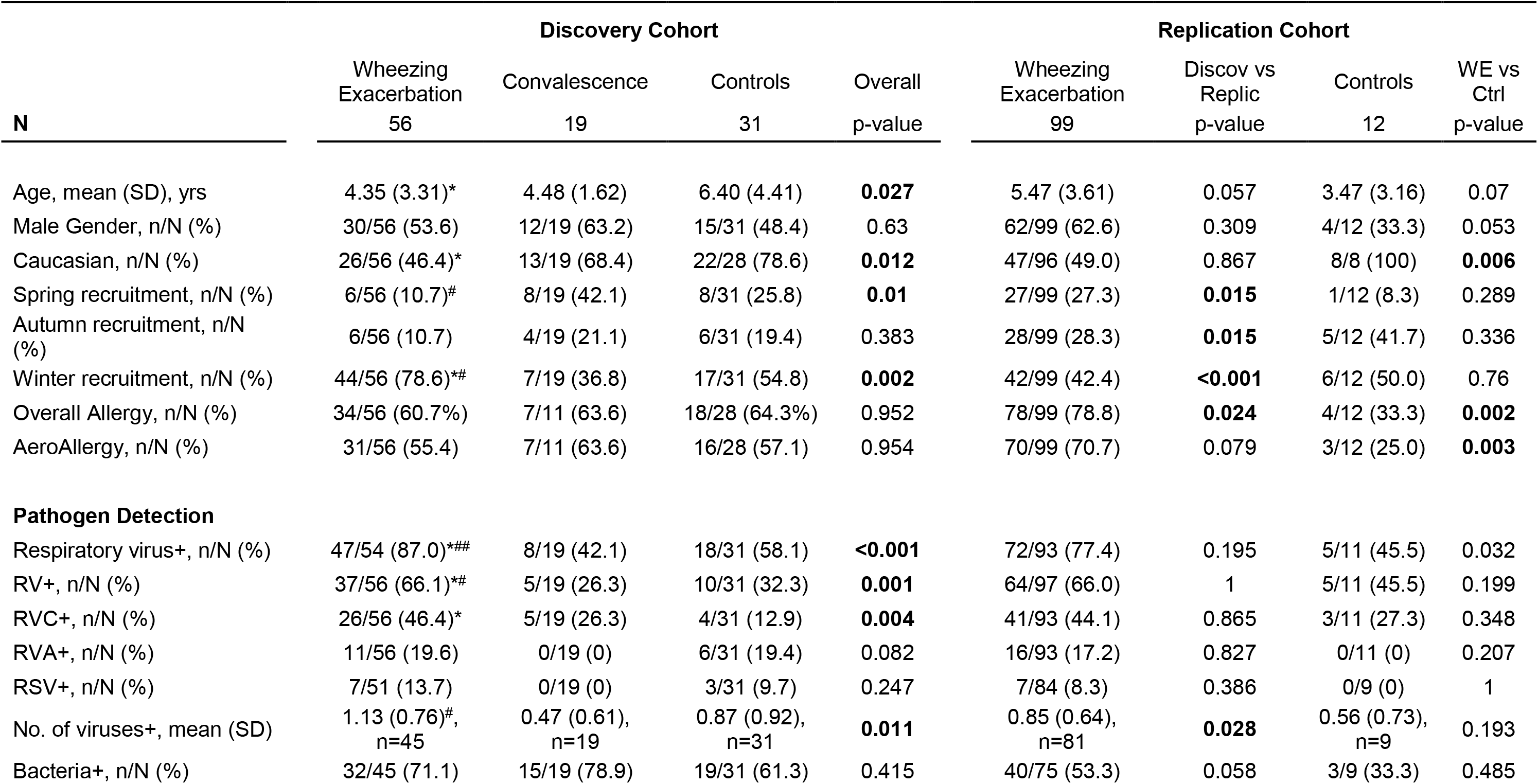
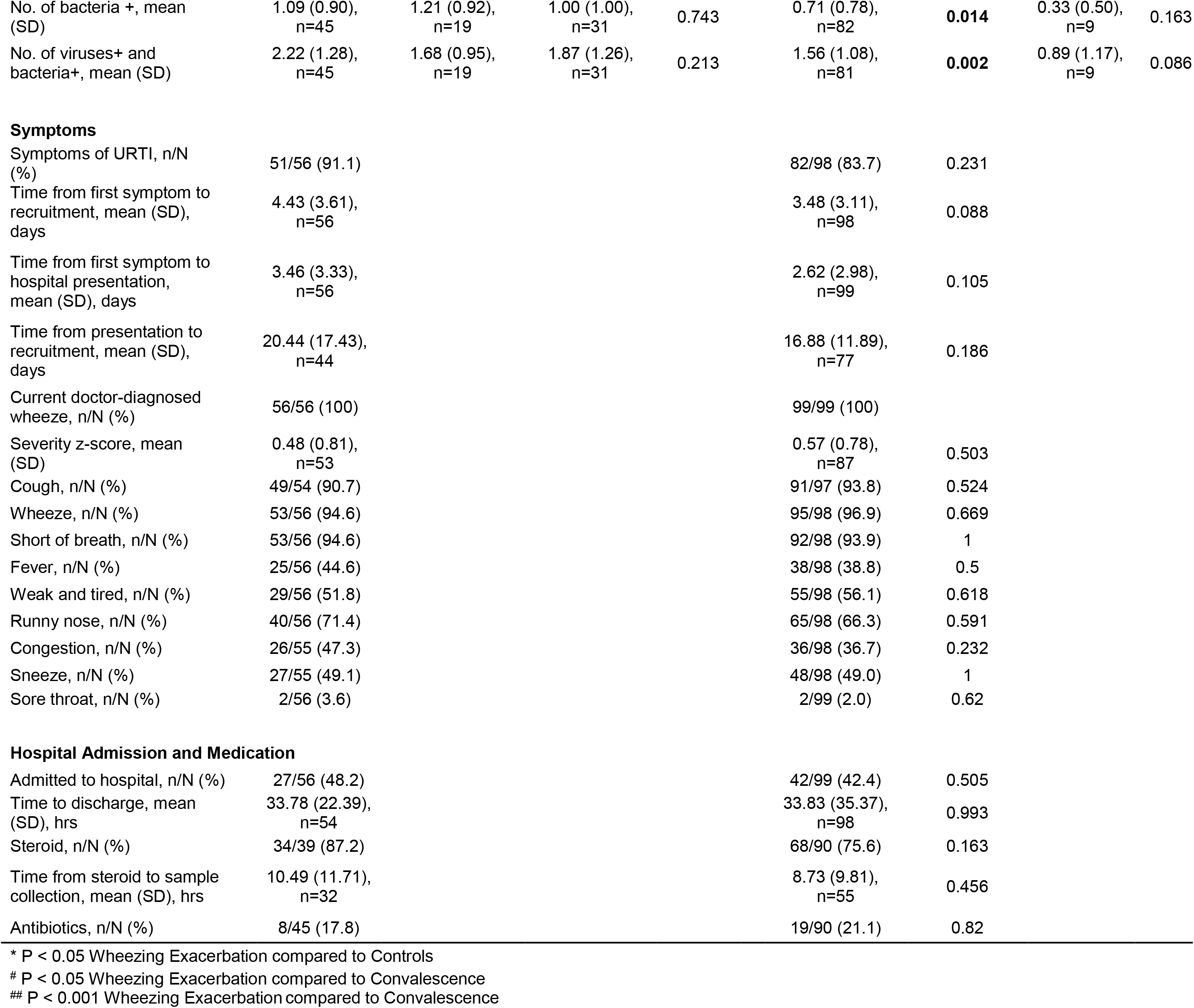
Characteristics of the study population.

Children with wheezing exacerbations were younger (mean = 4.35 (SD 3.31) years) than controls (6.40 (SD 4.41) years, p = 0.025) and had fewer Caucasians (46.4% vs 78.6%, p = 0.006). A higher proportion of the children with wheezing exacerbations were sampled during winter, compared to convalescence (p = 0.001) and controls (p = 0.028). Respiratory virus, in particular, RV, was widely detected during wheezing exacerbations (87.0%, and 66.1%, respectively) compared with convalescence (42.1% and 26.3%, p < 0.001 and p = 0.003, respectively) and controls (58.1% and 32.3%, p = 0.004 and p = 0.003, respectively). RV-C was more prevalent during wheezing exacerbations (46.4%) compared with controls (12.9%, p = 0.002). The number of viruses detected during acute exacerbations (1.13 (SD 0.76) was higher than during convalescence (0.47 (SD 0.61), p = 0.001). No difference in bacterial detection was observed between the groups.

The proportion of children with wheezing exacerbations recruited in winter in the replication cohort was lower than the discovery cohort (p < 0.001, Table 1). The replication cohort was also more atopic (78.8% vs 60.7%, p = 0.024), and had lower detection rates of respiratory viral and bacterial pathogens compared to the latter. Respiratory symptoms, hospital admissions and medication usage were not different in the discovery and replication cohorts.

### Gene expression profiling of exacerbation responses in the upper airways

Nasal swab specimens were collected from the children and gene expression patterns were profiled on microarrays. Cellular composition data of the samples was not available, and therefore a computational approach was employed to estimate the proportions of different cell types directly from the microarray profiles.^31^ As illustrated in Figure 1a, the highest proportions were observed for neutrophils, epithelial cells, and monocytes. Stratification of the subjects by case/control status and RV detection revealed that wheezing exacerbations were associated with increased proportions of M1 macrophages and decreased proportions of conventional dendritic cells (Figure 1b). The proportions of the other cell types were mostly comparable across the groups. The relationship between case/control status, RV detection and cellular composition for each individual child is illustrated in Figure 1c.

**Figure 1.**
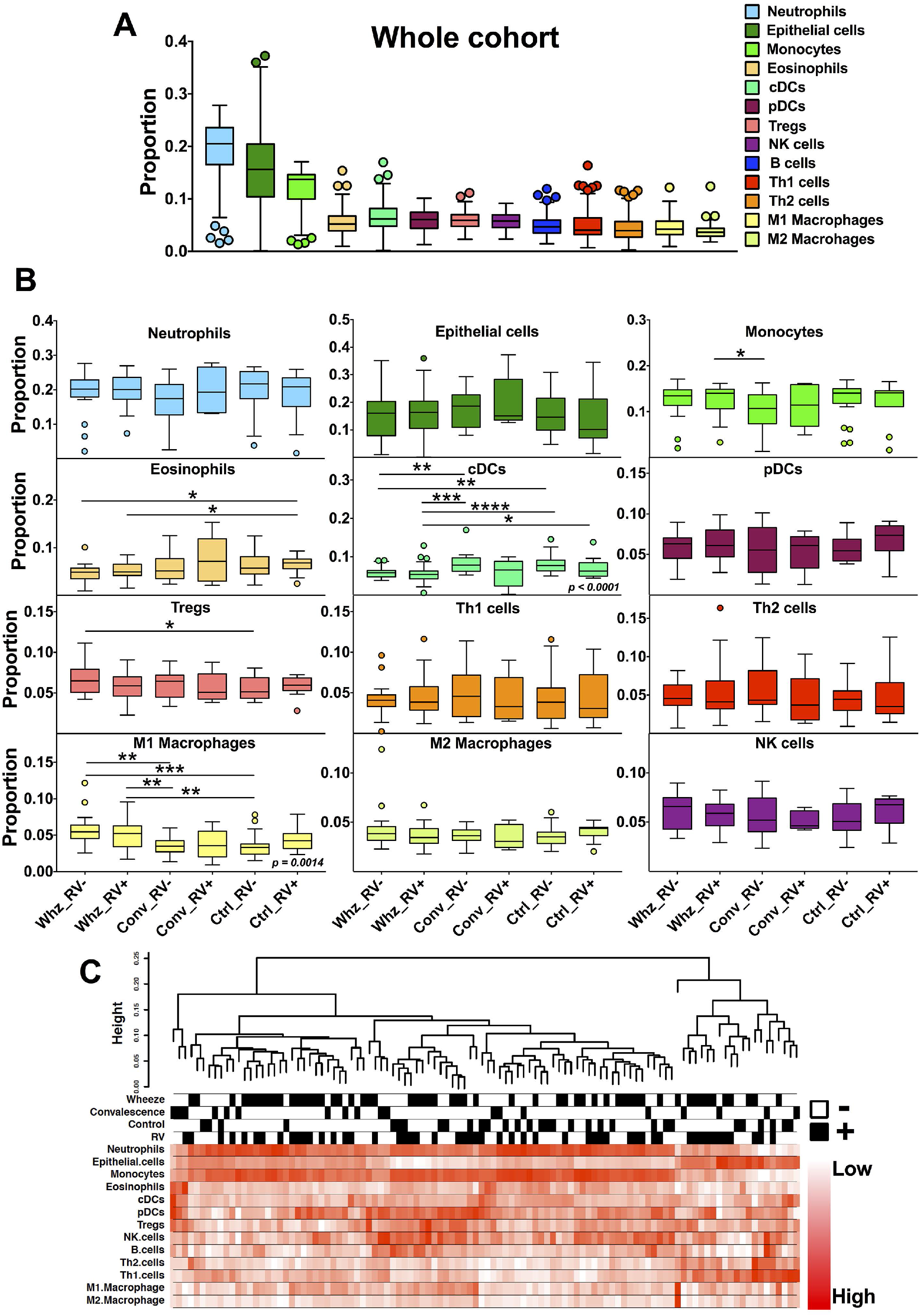
Computational inference of the cellular composition of the nasal swab samples. **A)** Relative proportions of 12 major cell types across the cohort. **B)** Relative proportions of individual cell types stratified according to RV status (RV+, RV-) within wheeze (Whz), convalescent (Conv) and control (Ctrl) subjects. **C)** Relationship between cell type proportions and clinical traits. Proportions were determined by computational deconvolution. Boxplots show IQR fenced using the Turkey method. Significant two-way Kruskal-Wallis P values are shown in italics. * represents a Mann-Whitney P value < 0.05, ** < 0.01, *** < 0.001, and **** < 0.0001.

To identify differentially expressed genes, we undertook group-wise comparisons employing *LIMMA* in combination with *Surrogate Variable Analysis*, which adjusts the analysis for all estimated sources of unwanted biological and technical variation (See Methods). Comparison of gene expression patterns between children with RV positive wheezing exacerbations (n=37) versus RV negative controls (n=21) revealed that 137 genes were upregulated and 82 genes were downregulated (adjusted p-value < 0.05, Table S1). Pathways analysis demonstrated that the upregulated genes were highly enriched for type I interferon signalling (adjusted p-value = 4.8 × 10^−22^, relevant genes are highlighted in Table S1). Type I interferon signalling (adjusted p-value = 8.5 × 10^−31^, Table S2) was also upregulated in children with RV positive exacerbations in comparison to RV negative convalescent children (n=14).

Gene expression patterns were then assessed in children with RV negative exacerbations. Comparison of these children (n=19) with RV negative controls (n=21) revealed that 17 genes were upregulated and 10 genes were downregulated (adjusted p-value < 0.05, Table S3). Similar results were obtained by comparing children with RV negative exacerbations versus RV negative convalescent children (n=14), where 30 differentially expressed genes were identified (adjusted p-value < 0.05, Table S4). While a subset of these genes were also upregulated in children with RV positive exacerbations (e.g. IL-18R1, CD163), a notable difference was the lack of an interferon signature.

### Discovery of molecular sub-phenotypes of acute wheezing illnesses

The findings above suggested that children with RV positive exacerbations mount an interferon response but there appears to be additional heterogeneity. As the data analysis strategy relied on group-wise comparisons (with LIMMA/SVA), this approach cannot shed any insight into subject-to-subject variations in gene expression patterns within each group. To obtain more detailed information in this regard, we employed hierarchical clustering (See Methods). To ensure that the clustering did not simply reflect variations in cellular composition (observed above in Figure 1), we utilised a set of negative control genes (i.e. genes not related to the outcome of interest) to model unwanted variation in the data, and removed these effects using regression (See Methods and Figure S1). This strategy preserves the gene expression signature associated with acute exacerbations in the data, and removes the strong correlation structure between samples that reflect variations in cellular composition (Figure S1). As illustrated in Figure 2a, cluster analysis of the corrected data divided the subjects into five distinct clusters or molecular phenotypes (labelled S1-S5). Likewise, the genes were also divided into five clusters labelled G1-G5. Notably, the majority of children with exacerbations (45/56 = 80%) were found within clusters S1 and S2, and there was no difference in RV detection between these cluster groups of children (RV detection rates for children with exacerbations in cluster S1 vs cluster S2 were 73% vs 58%, p-value = 0.347). The other three clusters S3-S5 contained the bulk of the controls and the convalescent samples, as well as 11 remaining children with exacerbations.

**Figure 2.**
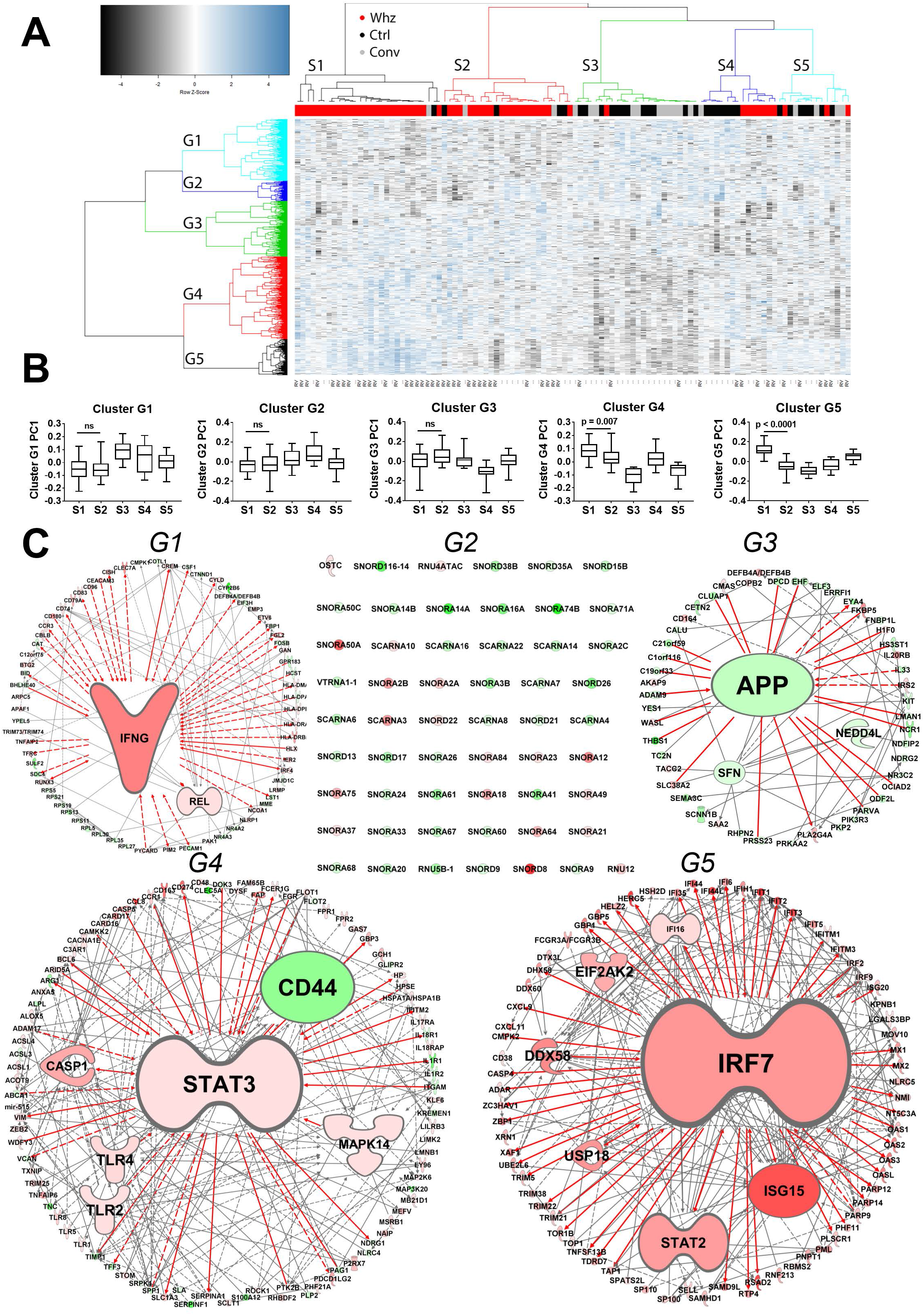
Discovery of IRF7 molecular phenotypes. **A)** Consensus hierarchical clustering was performed on gene expression profiles derived from nasal swab samples. Five clusters of subjects (S1-S5) and genes (G1-G5) were identified. **B)** The expression profiles of each gene cluster were summarized by principal components analysis, and the first principal component of each gene cluster was plotted across the subjects stratified by their cluster membership. **C)** Experimentally supported findings from published studies (prior knowledge) were employed to reconstruct the wiring diagram of the gene networks for each gene cluster. Genes colored red were upregulated and those colored green were downregulated in subjects from cluster S1 versus S2. Solid/dashed lines indicate direct/indirect functional relationships between genes. Larger nodes have more connections.

To examine the overall expression of each gene cluster across the children, principal component analysis was employed to summarise the expression data for each cluster, and the first principal component was plotted across the subjects, stratified into their respective clusters. This analysis revealed that the most striking difference between the subjects in cluster groups S1 and S2 was the differential expression of the set of genes in cluster G5 (Figure 2b), and this cluster of genes was strongly enriched for type I interferon signalling (data not shown). Moreover, prior knowledge based reconstruction of the wiring diagram of the underlying gene networks for each cluster revealed that IRF7 – a master regulator of type I interferon responses,^32^ was the dominant hub gene identified in cluster G5. Accordingly, we designated the children in clusters S1 and S2 as IRF7^hi^ versus IRF7^lo^ molecular phenotypes.

### Network hubs and driver genes underlying IRF7^hi^ versus IRF7^lo^ molecular phenotypes

To further characterize the IRF7 phenotypes, children with exacerbations were stratified into IRF7^hi^ (n=26) and IRF7^lo^ (n=19) subgroups, and compared with RV negative controls employing LIMMA/SVA. We followed this strategy because a direct comparison of IRF7^hi/low^ phenotypes would only reveal differences between the respective responses, and we wanted to know if the phenotypes operate through discrete and/or overlapping pathways. The data showed that 208 genes were upregulated and 157 genes were downregulated in children with IRF7^hi^ exacerbations compared with controls (Figure 3a, left panel; Table S5). IRF7^lo^ exacerbations were characterized by upregulation of 96 genes, and downregulation of 31 genes downregulated (Figure 3a, right panel; Table S6). These analyses revealed an overlapping response signature, which comprised 52 upregulated genes and 11 downregulated genes. Gene network analysis revealed that IRF7 gene networks were upregulated in the children with IRF7^hi^ exacerbations (Figure 3b, left panel). IRF7^lo^ exacerbations lacked an IRF7 signature, and instead were characterized by upregulation of Th2-associated pathways (e.g. IL-4R, FCER1G, ARG1) and downregulation of IFNg (Figure 3b, right panel). We also built a combined network to illustrate the unique and overlapping gene network patterns for each phenotype (Figure 4).

**Figure 3.**
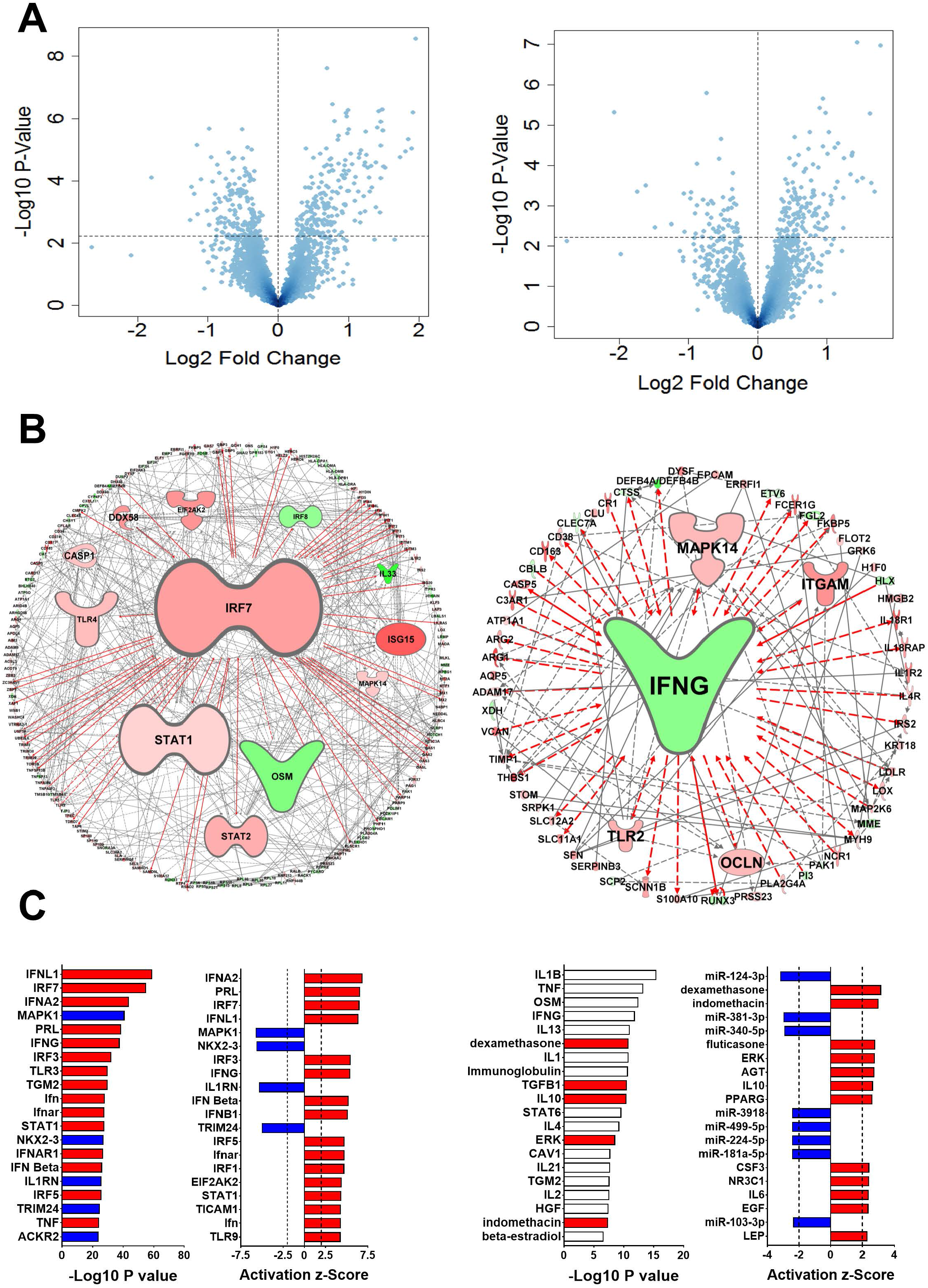
Identification of network hubs and driver genes underlying IRF7hi and IRF7lo exacerbations. **A)** Differentially expressed genes were identified by comparing children with IRF7hi exacerbations (left panel) or IRF7lo exacerbations (right panel) with RV negative controls. Data are presented as volcano plots, and the dashed horizontal line represents FDR = 0.05. **B)** The gene networks underlying IRF7hi exacerbations (left panel) or IRF7lo exacerbations (right panel) were reconstructed employing prior knowledge. Genes colored red were upregulated and genes colored green were downregulated. Larger nodes have more connections. **C)** Upstream regulator analysis was employed to infer molecular drivers of IRF7hi (left panel) and IRF7lo (right panel) exacerbations. The drivers were ranked by – Log10 overlap p-value or activation Z-score. Red bars indicate pathway activation (activation Z-score greater than 2.0) and blue bars indicate pathway inhibition (activation Z-score less than −2.0). White bars indicate pathways with non-significant activation Z-scores (i.e. absolute Z-scores less than 2.0).

**Figure 4:**
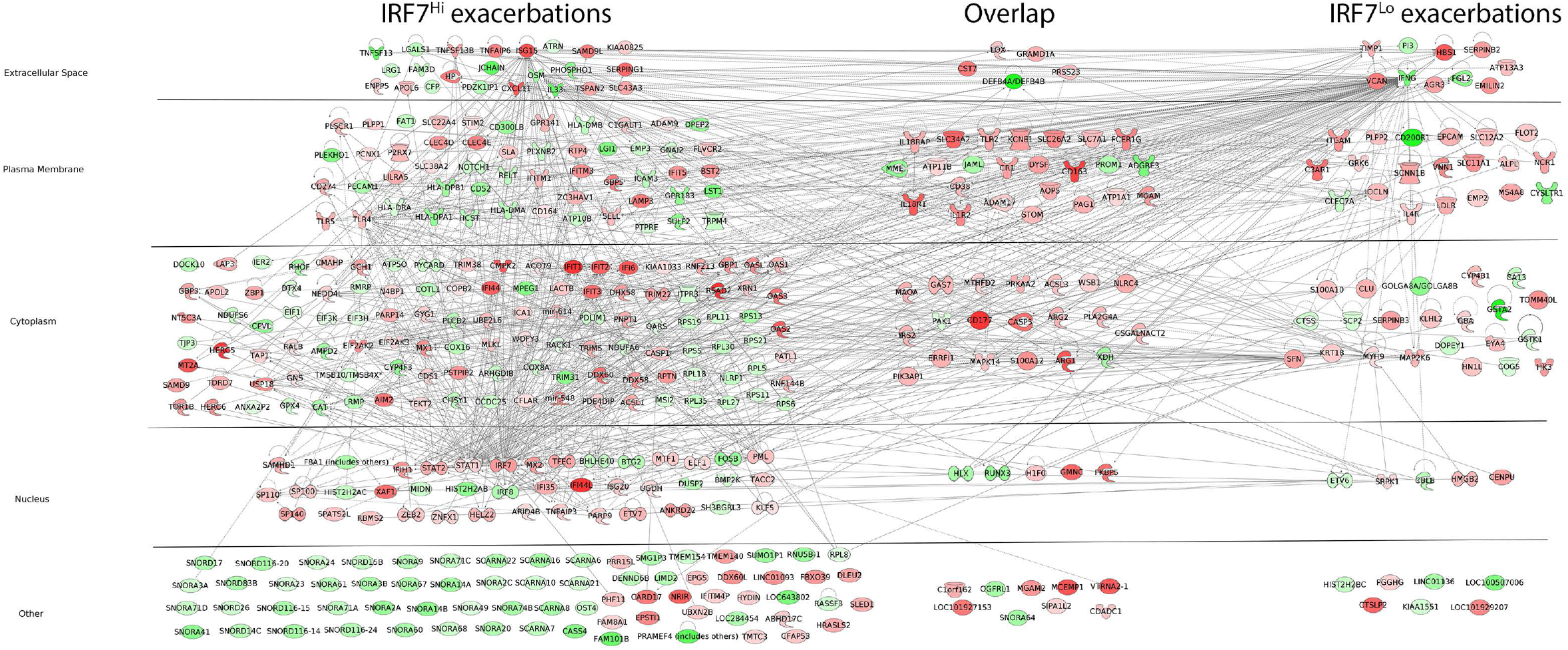
IRF7hi and IRF7lo exacerbation phenotypes operate through discrete and overlapping pathways. Gene networks associated with IRF7hi exacerbations (left side), IRF7lo exacerbations (right side), or common to both responses (middle). Genes are organised by subcellular location. Genes colored red were upregulated and those colored green were downregulated relative to RV negative controls.

Next we employed upstream regulator analysis to identify molecular drivers of IRF7^hi^ and IRF7^lo^ exacerbation responses.^28, 33^ The data showed that IRF7^hi^ exacerbations were putatively driven by upregulation of IFNL1, IRF7, and interferon alpha signalling (Figure 3c, left panel, Table S7). In contrast, IRF7^lo^ exacerbations were predicted to be driven by upregulation of cytokine and growth factor signalling pathways (e.g. IL-6, IL-10, TGFb, ERK, CSF3, EGF, Figure 3c, right panel, Table S8). It is noteworthy that multiple inflammatory pathways (e.g. IL-1b, IL-2, IL-4, IL-13, TNF, OSM, IFNG) featured in the upstream regulator analysis results for IRF7^lo^ exacerbations, however the activation Z-scores were not significant, indicating that the direction of the gene expression changes in terms of up- and down- regulation were not consistent with the known role of these pathways in the regulation of gene expression.^28^

We re-examined the cellular composition data from Figure 1 in IRF7^hi^ and IRF7^lo^ exacerbation responses, and we found that both phenotypes were associated with decreased proportions of cDCs, and similar proportions of M1 macrophages (Figure S2). Stratification of IRF7 phenotypes by RV detection revealed further heterogeneity, including upregulated proportions of Th2 cells in children with RV positive, IRF7^lo^ exacerbations (Figure S3).

### Candidate pathways linking exacerbation responses with asthma-related traits

To identify candidate pathways that potentially link exacerbation responses with expression of asthma-related traits, we employed Pubmatrix^34^ to screen the literature for relevant studies (Table S9). We found that multiple genes which were upregulated in children with IRF7^hi^ exacerbations (e.g. CASP1, CD274, CXCL11, DDX58, Haptoglobin, IFIH1, IRF7, P2RX7, PHF11, Selectin L, SERPING1, STAT1, TLR4, TLR5, TNFAIP3, TNFAIP6, TNFSF13B) or IRF7^lo^ exacerbations (e.g. AGR3, C3AR1, EPCAM, IL4R, LDLR, NCR1, OCLN, SERPINB2, SERPINB3, THBS1, TIMP1, VCAN) have been previously studied in the context of asthma and/or related traits. Moreover, overlap signature of genes that were commonly upregulated in children with IRF7^hi^ and IRF7^lo^ exacerbations have also been previously studied in this context (e.g. ADAM17, ARG1, ARG2, CD163, CD38, FCER1G, IL18R1, PLA2G4A, S100A12, TLR2).

### Clinical characteristics of IRF7^hi^ versus IRF7^lo^ exacerbations

We next investigated the biological and clinical characteristics of IRF7^hi^ and IRF7^lo^ exacerbations. These groups were not related to any technical variables measured in the study (Table S10). There was also no difference in season of recruitment, the detection of viral or bacterial pathogens, or the use of medications (Table 2). The prevalence of atopy, including allergy to aeroallergens, was higher in IRF7^hi^ exacerbations (76.9% and 73.1%, respectively) compared with IRF7^lo^ exacerbations (42.1% and 31.6%, p = 0.029 and 0.008, respectively). Children with IRF7^lo^ exacerbations presented to hospital much later after initial symptoms (4.74 (SD 4.03) days) than IRF7^hi^ exacerbations (2.31 (SD 1.98) days, p = 0.011). This was also reflected in the duration of cough prior to hospitalization (IRF7^lo^: 5.62 (SD 3.20) days and IRF7^hi^: 1.96 (SD 1.51) days, respectively, p = 0.000027). Children with IRF7^lo^ exacerbations were at least 4.6 times more likely to be admitted to hospital compared with IRF7^hi^ exacerbations (OR 4.65, p = 0.018). Time to subsequent first representation/admission to hospital with an exacerbation was shorter in IRF7^lo^ exacerbations compared with IRF7^hi^ exacerbations (Table 2, Fig S5). Within the first year of follow-up, more children with IRF7^lo^ (68.4%) represented/readmitted to hospital with a respiratory exacerbation compared to those with IRF7^hi^ (30.8%, p = 0.017). All associations remained significant after adjustment for age, gender, aeroallergen allergy and Caucasian ethnicity.

**Table 2.**
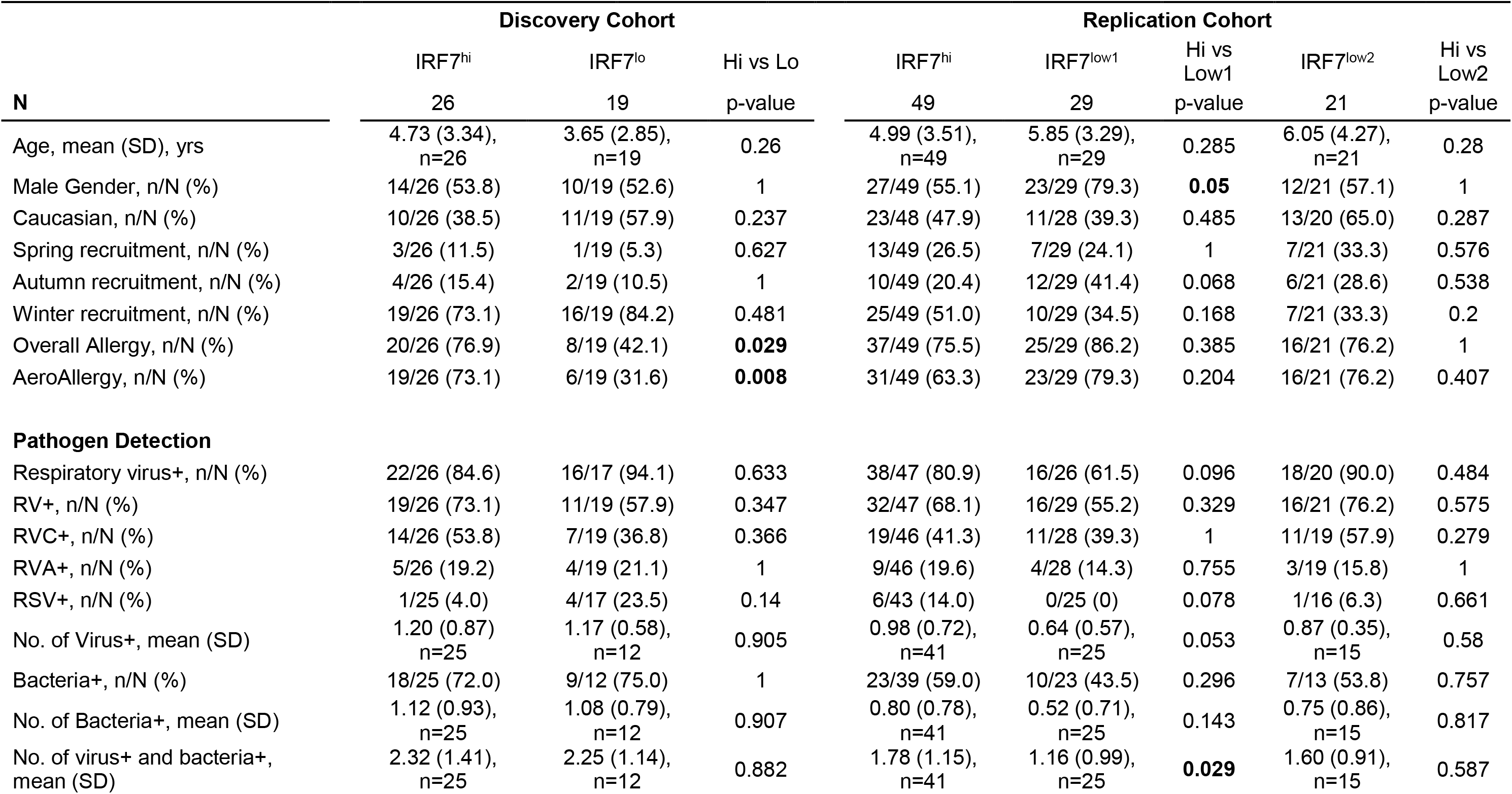
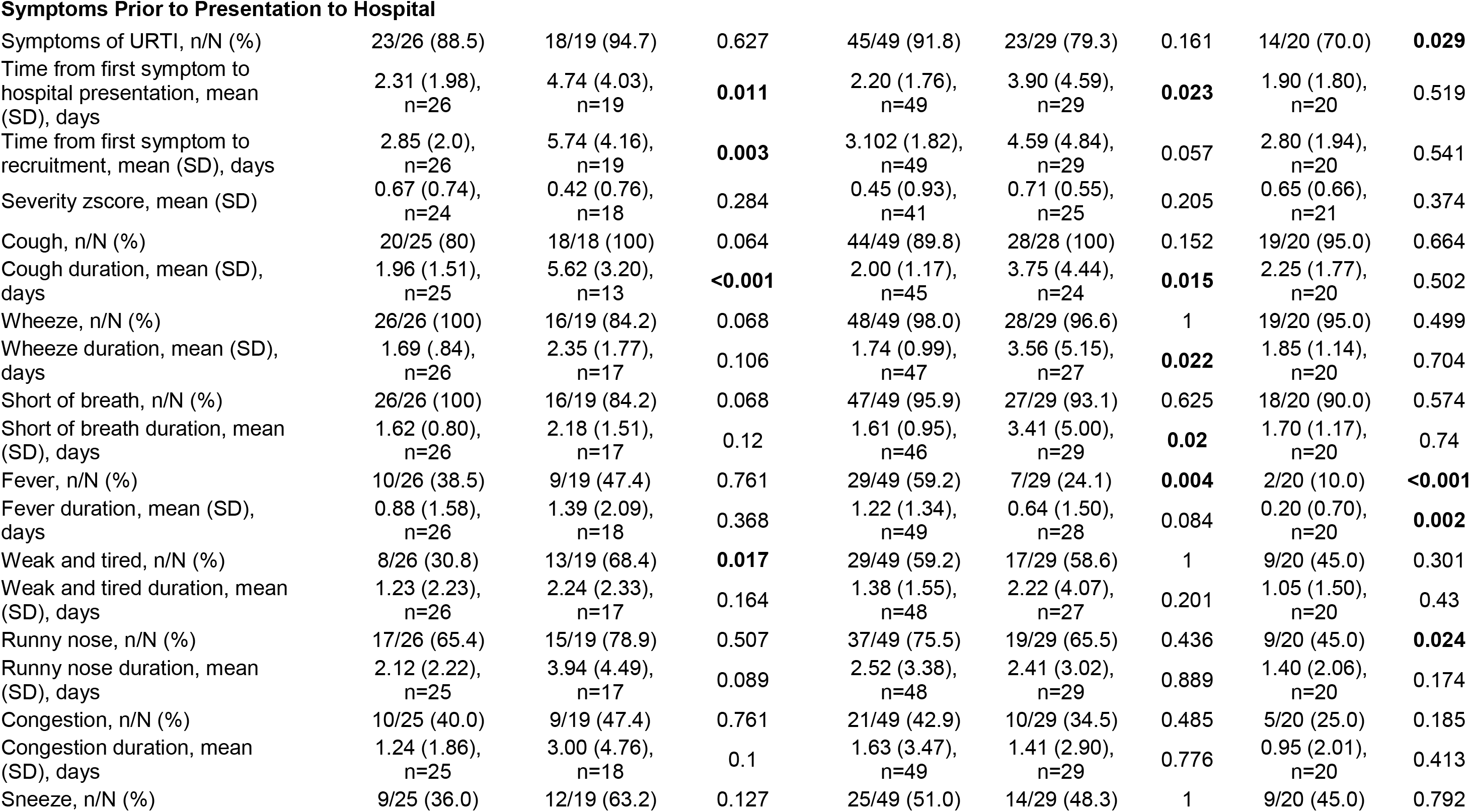
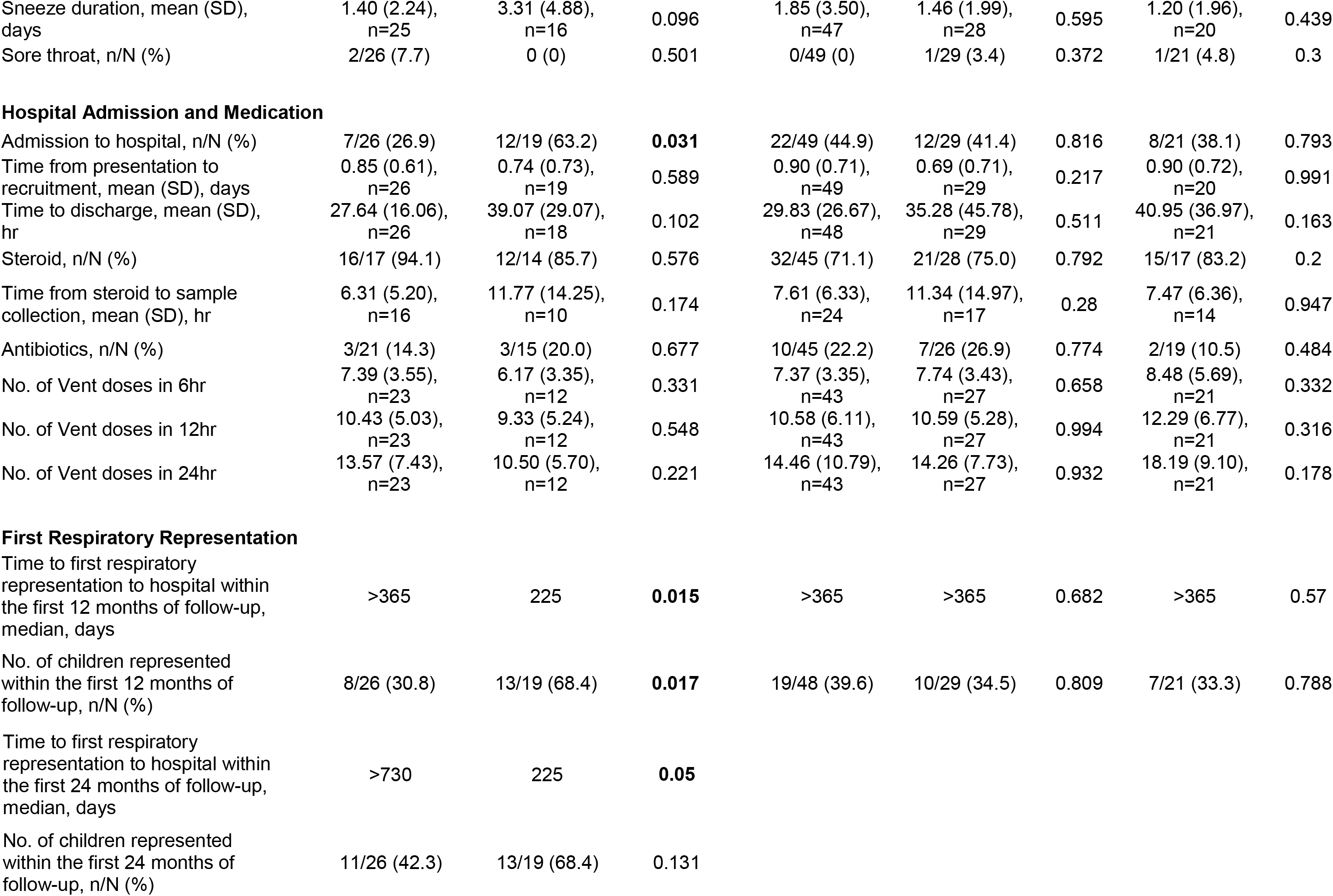
Characteristics of the IRF7^hi^ and IRF7^lo^ phenotypes in the discovery and replication cohorts.

| N | Discovery Cohort |  |  | Replication Cohort |  |  |  |  |
| --- | --- | --- | --- | --- | --- | --- | --- | --- |
|  | IRF7<sup>hi</sup> | IRF7<sup>lo</sup> | Hi vs Lo | IRF7<sup>hi</sup> | IRF7<sup>low1</sup> | Hi vs Low1 | IRF7<sup>low2</sup> | Hi vs Low2 |
|  | 26 | 19 | p-value | 49 | 29 | p-value | 21 | p-value |
| Age, mean (SD), yrs | 4.73 (3.34),<br>n=26 | 3.65 (2.85),<br>n=19 | 0.26 | 4.99 (3.51),<br>n=49 | 5.85 (3.29),<br>n=29 | 0.285 | 6.05 (4.27),<br>n=21 | 0.28 |
| Male Gender, n/N (%) | 14/26 (53.8) | 10/19 (52.6) | 1 | 27/49 (55.1) | 23/29 (79.3) | **0.05** | 12/21 (57.1) | 1 |
| Caucasian, n/N (%) | 10/26 (38.5) | 11/19 (57.9) | 0.237 | 23/48 (47.9) | 11/28 (39.3) | 0.485 | 13/20 (65.0) | 0.287 |
| Spring recruitment, n/N (%) | 3/26 (11.5) | 1/19 (5.3) | 0.627 | 13/49 (26.5) | 7/29 (24.1) | 1 | 7/21 (33.3) | 0.576 |
| Autumn recruitment, n/N (%) | 4/26 (15.4) | 2/19 (10.5) | 1 | 10/49 (20.4) | 12/29 (41.4) | 0.068 | 6/21 (28.6) | 0.538 |
| Winter recruitment, n/N (%) | 19/26 (73.1) | 16/19 (84.2) | 0.481 | 25/49 (51.0) | 10/29 (34.5) | 0.168 | 7/21 (33.3) | 0.2 |
| Overall Allergy, n/N (%) | 20/26 (76.9) | 8/19 (42.1) | **0.029** | 37/49 (75.5) | 25/29 (86.2) | 0.385 | 16/21 (76.2) | 1 |
| AeroAllergy, n/N (%) | 19/26 (73.1) | 6/19 (31.6) | **0.008** | 31/49 (63.3) | 23/29 (79.3) | 0.204 | 16/21 (76.2) | 0.407 |
| **Pathogen Detection** |  |  |  |  |  |  |  |  |
| Respiratory virus+, n/N (%) | 22/26 (84.6) | 16/17 (94.1) | 0.633 | 38/47 (80.9) | 16/26 (61.5) | 0.096 | 18/20 (90.0) | 0.484 |
| RV+, n/N (%) | 19/26 (73.1) | 11/19 (57.9) | 0.347 | 32/47 (68.1) | 16/29 (55.2) | 0.329 | 16/21 (76.2) | 0.575 |
| RVC+, n/N (%) | 14/26 (53.8) | 7/19 (36.8) | 0.366 | 19/46 (41.3) | 11/28 (39.3) | 1 | 11/19 (57.9) | 0.279 |
| RVA+, n/N (%) | 5/26 (19.2) | 4/19 (21.1) | 1 | 9/46 (19.6) | 4/28 (14.3) | 0.755 | 3/19 (15.8) | 1 |
| RSV+, n/N (%) | 1/25 (4.0) | 4/17 (23.5) | 0.14 | 6/43 (14.0) | 0/25 (0) | 0.078 | 1/16 (6.3) | 0.661 |
| No. of Virus+, mean (SD) | 1.20 (0.87)<br>n=25 | 1.17 (0.58),<br>n=12 | 0.905 | 0.98 (0.72),<br>n=41 | 0.64 (0.57),<br>n=25 | 0.053 | 0.87 (0.35),<br>n=15 | 0.58 |
| Bacteria+, n/N (%) | 18/25 (72.0) | 9/12 (75.0) | 1 | 23/39 (59.0) | 10/23 (43.5) | 0.296 | 7/13 (53.8) | 0.757 |
| No. of Bacteria+, mean (SD) | 1.12 (0.93),<br>n=25 | 1.08 (0.79),<br>n=12 | 0.907 | 0.80 (0.78),<br>n=41 | 0.52 (0.71),<br>n=25 | 0.143 | 0.75 (0.86),<br>n=15 | 0.817 |
| No. of virus+ and bacteria+, mean (SD) | 2.32 (1.41),<br>n=25 | 2.25 (1.14),<br>n=12 | 0.882 | 1.78 (1.15),<br>n=41 | 1.16 (0.99),<br>n=25 | **0.029** | 1.60 (0.91),<br>n=15 | 0.587 |
| Symptoms Prior to Presentation to Hospital |  |  |  |  |  |  |  |  |
| Symptoms of URTI, n/N (%) | 23/26 (88.5) | 18/19 (94.7) | 0.627 | 45/49 (91.8) | 23/29 (79.3) | 0.161 | 14/20 (70.0) | **0.029** |
| Time from first symptom to hospital presentation, mean (SD), days | 2.31 (1.98),<br>n=26 | 4.74 (4.03),<br>n=19 | **0.011** | 2.20 (1.76),<br>n=49 | 3.90 (4.59),<br>n=29 | **0.023** | 1.90 (1.80),<br>n=20 | 0.519 |
| Time from first symptom to recruitment, mean (SD), days | 2.85 (2.0),<br>n=26 | 5.74 (4.16),<br>n=19 | **0.003** | 3.102 (1.82),<br>n=49 | 4.59 (4.84),<br>n=29 | 0.057 | 2.80 (1.94),<br>n=20 | 0.541 |
| Severity zscore, mean (SD) | 0.67 (0.74),<br>n=24 | 0.42 (0.76),<br>n=18 | 0.284 | 0.45 (0.93),<br>n=41 | 0.71 (0.55),<br>n=25 | 0.205 | 0.65 (0.66),<br>n=21 | 0.374 |
| Cough, n/N (%) | 20/25 (80) | 18/18 (100) | 0.064 | 44/49 (89.8) | 28/28 (100) | 0.152 | 19/20 (95.0) | 0.664 |
| Cough duration, mean (SD), days | 1.96 (1.51),<br>n=25 | 5.62 (3.20),<br>n=13 | **<0.001** | 2.00 (1.17),<br>n=45 | 3.75 (4.44),<br>n=24 | **0.015** | 2.25 (1.77),<br>n=20 | 0.502 |
| Wheeze, n/N (%) | 26/26 (100) | 16/19 (84.2) | 0.068 | 48/49 (98.0) | 28/29 (96.6) | 1 | 19/20 (95.0) | 0.499 |
| Wheeze duration, mean (SD), days | 1.69 (.84),<br>n=26 | 2.35 (1.77),<br>n=17 | 0.106 | 1.74 (0.99),<br>n=47 | 3.56 (5.15),<br>n=27 | **0.022** | 1.85 (1.14),<br>n=20 | 0.704 |
| Short of breath, n/N (%) | 26/26 (100) | 16/19 (84.2) | 0.068 | 47/49 (95.9) | 27/29 (93.1) | 0.625 | 18/20 (90.0) | 0.574 |
| Short of breath duration, mean (SD), days | 1.62 (0.80),<br>n=26 | 2.18 (1.51),<br>n=17 | 0.12 | 1.61 (0.95),<br>n=46 | 3.41 (5.00),<br>n=29 | **0.02** | 1.70 (1.17),<br>n=20 | 0.74 |
| Fever, n/N (%) | 10/26 (38.5) | 9/19 (47.4) | 0.761 | 29/49 (59.2) | 7/29 (24.1) | **0.004** | 2/20 (10.0) | **<0.001** |
| Fever duration, mean (SD), days | 0.88 (1.58),<br>n=26 | 1.39 (2.09),<br>n=18 | 0.368 | 1.22 (1.34),<br>n=49 | 0.64 (1.50),<br>n=28 | 0.084 | 0.20 (0.70),<br>n=20 | **0.002** |
| Weak and tired, n/N (%) | 8/26 (30.8) | 13/19 (68.4) | **0.017** | 29/49 (59.2) | 17/29 (58.6) | 1 | 9/20 (45.0) | 0.301 |
| Weak and tired duration, mean (SD), days | 1.23 (2.23),<br>n=26 | 2.24 (2.33),<br>n=17 | 0.164 | 1.38 (1.55),<br>n=48 | 2.22 (4.07),<br>n=27 | 0.201 | 1.05 (1.50),<br>n=20 | 0.43 |
| Runny nose, n/N (%) | 17/26 (65.4) | 15/19 (78.9) | 0.507 | 37/49 (75.5) | 19/29 (65.5) | 0.436 | 9/20 (45.0) | **0.024** |
| Runny nose duration, mean (SD), days | 2.12 (2.22),<br>n=25 | 3.94 (4.49),<br>n=17 | 0.089 | 2.52 (3.38),<br>n=48 | 2.41 (3.02),<br>n=29 | 0.889 | 1.40 (2.06),<br>n=20 | 0.174 |
| Congestion, n/N (%) | 10/25 (40.0) | 9/19 (47.4) | 0.761 | 21/49 (42.9) | 10/29 (34.5) | 0.485 | 5/20 (25.0) | 0.185 |
| Congestion duration, mean (SD), days | 1.24 (1.86),<br>n=25 | 3.00 (4.76),<br>n=18 | 0.1 | 1.63 (3.47),<br>n=49 | 1.41 (2.90),<br>n=29 | 0.776 | 0.95 (2.01),<br>n=20 | 0.413 |
| Sneeze, n/N (%) | 9/25 (36.0) | 12/19 (63.2) | 0.127 | 25/49 (51.0) | 14/29 (48.3) | 1 | 9/20 (45.0) | 0.792 |
| Sneeze duration, mean (SD), days | 1.40 (2.24),<br>n=25 | 3.31 (4.88),<br>n=16 | 0.096 | 1.85 (3.50),<br>n=47 | 1.46 (1.99),<br>n=28 | 0.595 | 1.20 (1.96),<br>n=20 | 0.439 |
| Sore throat, n/N (%) | 2/26 (7.7) | 0 (0) | 0.501 | 0/49 (0) | 1/29 (3.4) | 0.372 | 1/21 (4.8) | 0.3 |
| Hospital Admission and Medication |  |  |  |  |  |  |  |  |
| Admission to hospital, n/N (%) | 7/26 (26.9) | 12/19 (63.2) | **0.031** | 22/49 (44.9) | 12/29 (41.4) | 0.816 | 8/21 (38.1) | 0.793 |
| Time from presentation to recruitment, mean (SD), days | 0.85 (0.61),<br>n=26 | 0.74 (0.73),<br>n=19 | 0.589 | 0.90 (0.71),<br>n=49 | 0.69 (0.71),<br>n=29 | 0.217 | 0.90 (0.72),<br>n=20 | 0.991 |
| Time to discharge, mean (SD), hr | 27.64 (16.06),<br>n=26 | 39.07 (29.07),<br>n=18 | 0.102 | 29.83 (26.67),<br>n=48 | 35.28 (45.78),<br>n=29 | 0.511 | 40.95 (36.97),<br>n=21 | 0.163 |
| Steroid, n/N (%) | 16/17 (94.1) | 12/14 (85.7) | 0.576 | 32/45 (71.1) | 21/28 (75.0) | 0.792 | 15/17 (83.2) | 0.2 |
| Time from steroid to sample collection, mean (SD), hr | 6.31 (5.20),<br>n=16 | 11.77 (14.25),<br>n=10 | 0.174 | 7.61 (6.33),<br>n=24 | 11.34 (14.97),<br>n=17 | 0.28 | 7.47 (6.36),<br>n=14 | 0.947 |
| Antibiotics, n/N (%) | 3/21 (14.3) | 3/15 (20.0) | 0.677 | 10/45 (22.2) | 7/26 (26.9) | 0.774 | 2/19 (10.5) | 0.484 |
| No. of Vent doses in 6hr | 7.39 (3.55),<br>n=23 | 6.17 (3.35),<br>n=12 | 0.331 | 7.37 (3.35),<br>n=43 | 7.74 (3.43),<br>n=27 | 0.658 | 8.48 (5.69),<br>n=21 | 0.332 |
| No. of Vent doses in 12hr | 10.43 (5.03),<br>n=23 | 9.33 (5.24),<br>n=12 | 0.548 | 10.58 (6.11),<br>n=43 | 10.59 (5.28),<br>n=27 | 0.994 | 12.29 (6.77),<br>n=21 | 0.316 |
| No. of Vent doses in 24hr | 13.57 (7.43),<br>n=23 | 10.50 (5.70),<br>n=12 | 0.221 | 14.46 (10.79),<br>n=43 | 14.26 (7.73),<br>n=27 | 0.932 | 18.19 (9.10),<br>n=21 | 0.178 |
| First Respiratory Representation |  |  |  |  |  |  |  |  |
| Time to first respiratory representation to hospital within the first 12 months of follow-up, median, days | >365 | 225 | **0.015** | >365 | >365 | 0.682 | >365 | 0.57 |
| No. of children represented within the first 12 months of follow-up, n/N (%) | 8/26 (30.8) | 13/19 (68.4) | **0.017** | 19/48 (39.6) | 10/29 (34.5) | 0.809 | 7/21 (33.3) | 0.788 |

|  |  |  |  |
| --- | --- | --- | --- |
| Time to first respiratory representation to hospital within the first 24 months of follow-up, median, days | >730 | 225 | **0.05** |
| No. of children represented within the first 24 months of follow-up, n/N (%) | 11/26 (42.3) | 13/19 (68.4) | 0.131 |

### Replication of IRF7^hi^ and IRF7^low^ exacerbation responses in an independent sample of children

To determine if IRF7^hi^ and IRF7^lo^ exacerbation phenotypes and their clinical correlates could be found in another group of children, RT-qPCR was employed to measure gene expression patterns in nasal swab samples from an independent sample of children. To select a panel of genes for RT-qPCR analysis, we focused on genes that were representative of IRF7^hi^ exacerbations (DDX60, IFNL1, IRF7, ISG15, Mx1, RSAD2, and the downregulated gene IL-33, Figure 4), IRF7^lo^ exacerbations (NCR1, THBS1, Figure 4), or that were common to both phenotypes (ARG1, CD163, FCER1G, IL-1R2, IL-18R1, TLR2, Figure 4). The RT-qPCR data was normalised to three endogenous control genes (HMBS, PPIA, PPIB), creating three separate variables for each gene, which were utilised for consensus clustering. The analysis segregated the subjects into five clusters (Figure 5); three of these had elevated expression of IRF7/interferon responses (combined as IRF7^hi^; black dendrogram in Figure 5), and the remaining two clusters had low IRF7/interferon responses (IRF7^low1^ and IRF7^low2^; red and green dendrograms respectively in Figure 5). A longer time lag was observed from first symptoms to hospital presentation in the IRF7^low1^ subjects (3.90 (SD 4.59) days) compared with the IRF7^hi^ group (2.20 (SD 1.76) days, p = 0.023). These symptoms included cough (p=0.015), wheeze (p=0.022) and shortness of breath (p = 0.02). Fever was more prevalent in the IRF7^hi^ group (59.2%) compared with the IRF7^low1^ and IRF7^low2^ groups (24.1%, p = 0.004 and 10.0%, p < 0.001, respectively). Runny nose was also more prevalent in the IRF7^hi^ group (75.5%) compared to the IRF7^low2^ group (45.0%, p = 0.024). All associations remained significant after adjusting for age, gender, aeroallergy and Caucasian ethnicity.

**Figure 5.**
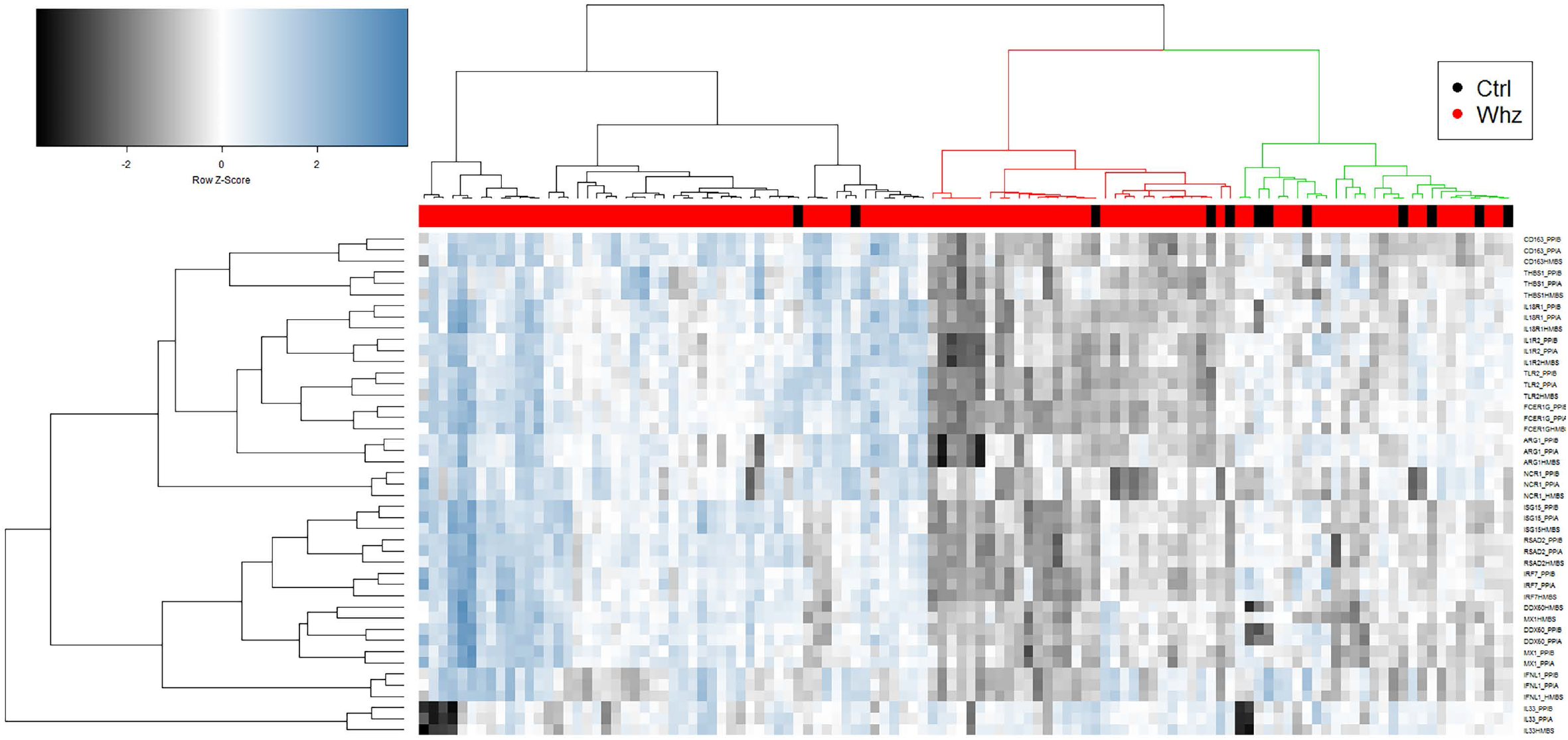
Identification of IRF7hi and IRF7lo exacerbations phenotypes in an independent cohort of children. Gene expression patterns were profiled by RT-qPCR in nasal swab samples collected from children with an asthma/wheezing exacerbation (red columns) or healthy controls (black columns). The RT-qPCR data was normalised separately to three independent endogenous reference genes, resulting in three variables for each target gene, and then analysed by consensus hierarchical cluster analysis. The dendrogram is colored by phenotype (IRF7hi = black, IRF7low1 green, IRF7low1 red).

## Discussion

Our study is the first to investigate acute wheeze/asthma exacerbation phenotypes in children using a systems biology approach. We have confirmed the central role of IRF7 as a gene network hub and have identified two distinct IRF7 molecular phenotypes, IRF7^hi^ exhibiting robust upregulation of the Th1/type I interferon response and IRF7^lo^ with an alternative activation signature marked by upregulation of cytokine and growth factor signalling and downregulation of interferon gamma. Importantly, the two phenotypes also produced distinct clinical phenotypes. Compared with children with IRF7^hi^, those with IRF7^lo^ had a slower progression of illness from initial symptoms to hospital presentation, a greater likelihood of a hospital admission, and a greater chance of representation with a further exacerbation. Exacerbation severity at presentation was not different between the two IRF7 patterns, perhaps because presentation to hospital is likely to be determined by symptoms reaching a severity threshold that causes parental concern. However also worth noting is that the apparently impaired IRF7 response in IRF7^lo^ children was not associated with an increase in exacerbation severity at presentation, although perhaps expectedly the reduced IRF7 response was associated with slower resolution of the episode. These findings were unaffected by the respiratory viruses or bacteria detected and medication use. Investigation of a replication cohort using RT-qPCR produced similar findings despite variations in the subject characteristics between the discovery and replication cohorts. In summary our findings reveal two distinct phenotypes with clear differences in gene regulation patterns, either upregulation of robust innate immune responses or cytokine and growth factor signalling, and associated differences in clinical characteristics.

Asthma and wheezing exacerbations are largely triggered by RV infections but the underlying mechanisms are not well understood.^2^ Previous *in vitro* studies suggested that RV-induced interferon responses were deficient in bronchial epithelial cells from subjects with asthma, resulting in increased viral loads and exaggerated secondary responses.^3, 4, 35^ However, *in vivo* studies of immune response patterns in the airways of both children and adults with RV-induced exacerbations found that interferon responses were increased, not deficient.^9–11, 36^ Moreover, a clinical trial that evaluated the utility of inhaled interferon beta therapy in adults at the first signs of a cold to prevent worsening of their asthma symptoms failed to achieve its primary endpoint.^37^ Our data extend these previous findings by demonstrating that wheezing exacerbations in children are heterogeneous and characterised by two very different IRF7 molecular phenotypes. Upstream regulator analysis suggested that IRF7^hi^ exacerbations were driven by upregulation of IFNL1, IRF7, and interferon alpha signalling. In contrast, IRF7^lo^ exacerbations were putatively driven by upregulation of cytokine and growth factor signalling (i.e. IL-6, IL-10, TGFb, CSF3, EGF) and downregulation of interferon gamma.

IRF7 is a master regulator of type I and type III interferon gene expression.^32, 38–40^ We previously reported that IRF7 gene networks were upregulated in nasal wash samples from asthmatic children experiencing mild-to-moderate viral exacerbations.^10^ We have also shown that IRF7 promotes RV-induced innate antiviral responses and limits IL-33 mRNA expression in cultured bronchial epithelial cells,^33^ and furthermore in our current study IL-33 was downregulated in IRF7^hi^ exacerbation responses. Girkin *et al*. reported that knockdown of IRF7 abolished RV-induced type I interferon responses in the airways in a mouse model.^40^ Together, these experimental findings support our computational analysis unveiling IRF7 as a regulator of the gene networks underlying the IRF7^hi^ responder phenotype.

Children with IRF7^lo^ exacerbations in the discovery cohort exhibited a delayed progression from first symptom onset to hospital presentation. Given that our study design entailed recruitment of children at ED presentation, we could not determine if IRF7 gene networks were initially upregulated closer to the onset of infection and subsequently waned, or alternatively if they were never upregulated in the first place. To address this issue, an alternative study design would be required which entails regular sampling of exacerbation-prone children during the RV season.^41^ It would also be of interest to further study the stability of these identified phenotypes to learn if subjects experience the same type of response over multiple wheezing exacerbation events or not. Notwithstanding this, our replication cohort comprised approximately twice as many cases as the discovery cohort, and this larger sample enabled the identification and characterization of two subgroups of IRF7^lo^ children, and only one of these subgroups was characterized by delayed progression. The activation of growth factor signalling pathways and downregulation of interferon gamma may reflect the immune response entering a resolution phase, however, at the same time these children were symptomatic. Moreover, many were also RV positive, and given that TGFb signalling promotes rhinovirus replication, upregulation of this pathway may prolong infection and delay viral clearance.^42, 43^ It is also known that frequent severe exacerbations are associated with deficits in lung function growth (children) and accelerated lung function decline (adults).^44^ Thus repeated cycles of inflammation, growth factor signalling and repair may alter the structure and function of the airways underlying this phenotype.

The mechanisms that determine expression of disease symptoms amongst children with IRF7^hi^ and IRF7^lo^ exacerbations are unknown. Given that children with IRF7^hi^ exacerbations elicit robust antiviral responses, one possibility is that the airways of these children are sensitive to the host response to respiratory viruses. In this regard, several pathways associated with the IRF7^hi^ phenotype are known to impact on respiratory function. PD-L1 (encoded by CD274) is an immune checkpoint that delivers an inhibitory signal for T cell activation. Upregulation of PD-L1 during respiratory bacterial infections in early life suppresses the IL-13 decoy receptor IL-13Ra2, resulting in persistent airways hyperresponsiveness.^45^ MDA5 (encoded by IFIH1) is a pattern recognition receptor that senses RV-derived RNA. MDA5 deficient mice infected with RV have delayed type I interferon responses, impaired type III interferon responses, and reduced airways hyperresponsiveness.^46^ The proinflammatory effectors TNF, IFNg, and interferon gamma-induced protein 10 can also promote airway hyper-responsiveness in animal models.^47–49^

Another possibility is that IRF7^hi^ and IRF7^lo^ exacerbation responses converge on a final common pathway to precipitate respiratory symptoms (Figure 4). For example, CD38 is a receptor with enzymatic activity, which hydrolyses NAD, generating reaction products that modulate calcium signalling. It is expressed on immune and airway smooth muscle cells, and it plays a dual role in asthma by enhancing airways inflammation and contractile responses in smooth muscle.^50^ FCER1G is a component of the high affinity IgE receptor. Anti-IgE therapy neutralises serum IgE, reduces the expression of the high affinity IgE receptor on dendritic cells and mast cells, and it also reduces the frequency of asthma exacerbations.^51, 52^ Phospholipases A2 are involved in the generation of eicosanoids from arachidonic acid. Knock-in of human sPLA2 into mice enhances airways inflammation and airways hyperresponsiveness.^53^ TLR2 is a pattern recognition receptor that acts as a sensor for RV capsid.^54^ In a mouse model of combined RV and allergen exposure, TLR2 signaling in macrophages was required for induction of airways inflammation and airways hyperresponsiveness.^55^ In summary, our data has identified multiple candidate pathways that link IRF7^hi^ and IRF7^lo^ exacerbation responses with expression of respiratory symptoms, and these pathways represent logical candidates for future drug development programs.

This study has limitations that should be acknowledged. The expression profiles were derived from nasal swab samples that comprised a mixed cell population. Follow-up studies employing focused analyses in individually isolated cell types or single cell transcriptomics will be required to dissect the role of specific subpopulations of cells in this disease. The study participants were sampled during natural exacerbations, and it is not possible to control for all of the variables that may potentially impact on the data (e.g. age, gender, ethnicity, natural allergen exposure, pathogen strains and combinations, medications). To address this issue, our analysis strategy employed surrogate variable analysis to systematically estimate and adjust the analysis for all sources of hidden biological and/or technical variation. Some of the clinical characteristics associated with IRF7 phenotypes in the discovery cohort did not replicate in the validation cohort. This may be due in part to variations in the demographics and clinical characteristics between the two cohorts, as well as the fact that IRF7 phenotypes were defined in the discovery cohort by microarray analysis of a large number of genes, whereas in the validation cohort it was based on RT-qPCR analysis of a restricted gene panel. Finally, while our analyses have characterised gene network patterns underlying exacerbation phenotypes and unveiled candidate molecular drivers of the responses, further studies will be required to dissect the mechanisms that give rise to these phenotypes and drive the expression of respiratory symptoms. Notwithstanding these limitations, our findings demonstrate that exacerbation responses in children are heterogeneous and comprise IRF7^hi^ versus IRF7^lo^ molecular phenotypes that determine clinical phenotypes. Future clinical trials targeting the interferon system in this disease should be stratified on the basis of these IRF7 phenotypes.

## Author Contributions

Conception and design of the study: PNS, IAL, AB; Acquisition of data: SKK, KF, JB, NT, FP, JE, SO, MB; Data analysis and interpretation: SKK, JR, GZ, LC, CM, LO, NT, RM, PNS, IAL, AB; Drafting the manuscript for important intellectual content: SKK, PNS, IAL, AB; all authors reviewed and approved the final manuscript.

## Acknowledgements

We thank all the children and their families who agreed to take part in the study. The study was funded by National Health and Medical Research Council (NHMRC APP1045760) and AstraZeneca. JB was funded by a Thoracic Society of Australia and New Zealand/AstraZeneca Respiratory Research Fellowship. LC is a recipient of the Asthma Foundation Fiona Staniforth and Australian Government Research Training Program Scholarships. NT is funded by the Mickey Hardy Asthma Australia National Scholarship. AB is supported by Fellowships from the Simon Lee Foundation and the Brightspark and McCusker Foundations.

## Method

### Study Participants

The participants were part of an ongoing study examining the mechanisms of acute viral respiratory infection in children (MAVRIC). Cases were children aged 0-16 years presenting to the ED of a tertiary children’s hospital (Princess Margaret Hospital, Perth, Western Australia) with acute wheeze and the availability of nasal fluid and swab specimens. All respiratory diagnoses were determined by the treating physician and were independent of study staff. Controls consisted of siblings of the cases or randomly selected children from the community. We defined ‘admission to hospital’ as admission to a non-ED hospital ward. We also collected 19 convalescent/quiescent nasal samples from children who were followed-up at least 6 weeks after an acute exacerbation of asthma or wheeze; only a subset of these samples (5/19) were paired with acute samples. The hospital’s Human Ethics Committee approved the study (MAVRIC approval 1761 EP) and parental/guardian written informed consent was obtained prior to recruitment. An independent sample of children from within the MAVRIC cohort was used as a validation cohort.

### Data and sample collection

A detailed questionnaire and medical records were used to provide demographic and medical information for each participant during recruitment and follow-up. A child was considered positive for aeroallergy if they had a positive response to either (1) the skin prick test completed on 9 allergens (rye grass, mixed grasses, dog, cat, cockroach, *Alternaria tenuis, Aspergillus fumigatus, Dermatophagoides farinae, D. pteronyssinus*), or (2) a positive specific IgE to either cat or house dust mite (>0.35kU/L) at either the acute or convalescent visit, or positive answers to the acute questionnaire for the questions “Does your child suffer from hayfever?”, or “Does your child suffer from allergies to: (a) grasses/pollens?; or (b) dust mite?”. A child was considered positive for allergy overall if they had (1) aeroallergy; or (2) a positive response to the skin prick tests for either cows’ milk or egg white, or (3) a parental report of a history of anaphylaxis, or (4) a high total IgE at either the acute or convalescent visit.

Acute asthma severity scores were assigned to each child at recruitment using a modified National Institute of Health score for children over 2 years of age^12^ and included assessments of respiratory rate appropriate for age, oxygen saturation, auscultation, retractions and dyspnea. A separate, age-appropriate severity score was used for children aged under 2 years^13^ with assessments of heart rate, respiratory rate, wheezing and accessory muscle use. Separate severity Z scores were calculated for each child within each of the two age groups to provide standardized scores across the whole cohort.

The time to next re-presentation or admission to a public hospital in Western Australian with any acute respiratory illness for each child, within the first and second year of observation, was obtained from the state hospital database retrospectively.

A nasal secretion sample from each participant was collected to test for the presence of respiratory viruses and bacteria. Nasal swab specimens were obtained from each child using flocked swabs (Copan, Italy) and were taken immediately to the laboratory for processing for gene expression profiling.

### Virus and bacteria detection

Common respiratory pathogens (adenovirus, respiratory syncytial virus (RSV) types A and B, bocavirus, coronavirus, parainfluenza viruses 1-4, influenza viruses A, B and C, metapneumovirus, *Bordetella* species, *Mycoplasma pneumoniae, Chlamydophila pneumoniae, Haemophilus influenzae, Pneumocystis jirovecii, Staphylococcus aureus, Streptococcus Pneumoniae* and *pyogenes*) were identified using a tandem multiplex real-time PCR assay as previously described.^14^ Rhinovirus (RV) was detected and genotyped by a molecular method as previously described.^15, 16^

### Microarray analysis of gene expression

Total RNA was extracted from nasal swabs using TRIzol (Invitrogen) followed by RNeasy MinElute (QIAgen).^10^ The quality and integrity of the RNA was analysed on the nanodrop and bioanalyzer (Agilent). Total RNA samples were shipped on dry ice to the Ramaciotti Centre for Genomics, at the University of New South Wales, Sydney, for processing and hybridization to Human Gene 2.1 ST microarrays (Affymetrix). The raw microarray data are available from the gene expression omnibus repository (accession: GSE103166).

### Microarray data analysis

The microarray data was preprocessed in R employing the Robust Multi-Array (RMA) algorithm.^17^ A custom chip description file (hugene21sthsentrezg; version 20) was utilized to map probe sets to genes.^18^ The quality of the microarray data was assessed employing the R package arrayQualityMetrics. Low quality/outlying arrays were identified on the basis of Relative Log Expression (RLE) and Normalized Unscaled Standard Error (NUSE) metrics and removed from the analysis.^19, 20^ Differentially expressed genes were identified using Linear Models for Microarray Data (LIMMA) in conjunction with Surrogate Variable Analysis (SVA).^21, 22^ Limma entails fitting a linear model to the log expression intensities for each gene, and Empirical Bayes methods are employed to moderate the standard error estimates resulting in improved power and more stable inference.^21^ Surrogate Variable Analysis estimates and captures all sources of hidden biological and/or technical variation that may potentially confound the analysis, and the estimated surrogate variables are added as covariates to the limma models. Where applicable, the correlation between paired samples was accounted for in the linear model fit through use of the duplicateCorrelation function. P-values derived from the limma models were adjusted for multiple testing employing the false discovery rate method.^21^ Non-informative probe sets were identified using the proportion of variation accounted by the first principal component (PVAC) algorithm and filtered out of the results from the differential expression analysis.^23^

### Discovery of molecular phenotypes

Molecular phenotypes were identified using consensus hierarchical clustering.^24^ This algorithm employs data resampling techniques to find consensus across thousands of runs of a cluster analysis.^24^ Prior to cluster analysis, batch-like effects and other sources of unwanted variation were modelled and removed using the RUVnormalize algorithm.^25^ This algorithm leverages a set of negative control genes that are not associated with the outcome of interest to model unwanted variation and regress it out of the data. To select negative control genes, genes associated with wheezing exacerbations were first identified in case/control comparisons between wheezing/convalescent or wheezing/control samples using LIMMA/SVA, and those genes with an adjusted p-value less than 0.2 for any comparison were excluded from control gene selection. Negative control genes (n=1000) were selected from the remaining genes using the empNegativeControls function from the RUVCorr package.^26^ After removal of unwanted variation using the RUVnormalize algorithm, the gene expression data was converted to Z-scores, and consensus hierarchical clustering was performed using the set of genes that were associated with the outcome of interest (adjusted p-value < 0.1). Hierarchical cluster analysis was performed using Pearson correlation and ward linkage.

### Pathways analysis

Pathways analysis was performed using Enrichr software and the Reactome database.^27^

### Prior knowledge based reconstruction of gene networks

Gene networks were constructed employing experimentally supported findings from published studies curated in the Ingenuity Systems Knowledge Base (Ingenuity Systems, Redwood City, California). Direct and indirect molecular relationships were considered from all available categories, including activation, inhibition, localisation, modification, molecular cleavage, phosphorylation, protein-DNA interaction, protein-protein interaction, regulation of binding, transcription, translation, and ubiquitination. The circular layout was employed to display the network graph object, and hubs were positioned at the centre of the network. The nodes were coloured based on the expression ratio; red nodes denote upregulated genes and green nodes denote downregulation.

### Upstream regulator analysis

Upstream regulator analysis (Ingenuity Systems, Redwood City, California) was employed to infer the molecular drivers of the observed differential gene expression patterns.^28^ Two statistical metrics were calculated. The overlap p-value measures enrichment of known downstream target genes for a given upstream regulator amongst the differentially expressed genes. The activation Z-score is a measure of the pattern match between the direction of the observed gene expression changes (in terms of up/down regulation) and the predicted pattern based on prior experimental evidence. Pathways with an activation Z-score greater than 2.0 are predicted to be activated, and an activation Z-score less than −2.0 indicates pathway inhibition. If the activation Z-score lies between −2.0 and 2.0, the activation state of the pathway cannot be predicted.

### Estimation of cellular composition from gene expression profiles

Cell type enrichment was examined in microarray-based gene expression profiles using the xCell webtool (University of California, San Francisco).^29^ Gene signatures for 64 cell types were determined from >1,800 transcriptomic profiles of purified cell samples allowing reliable enrichment analysis to investigate tissue heterogeneity. Cell types were chosen for further analysis based on their potential importance in the upper airways. Enrichment scores were converted to proportional cellular composition, allowed for by the linearity assumption, so that the sum of the proportions was equal to 1. Statistical comparisons were performed using nonparametric Kruskal-Wallis and Mann-Whitney U tests (GraphPad software [La Jolla, USA]).

### RT-qPCR

Total RNA extracted from nasal swab samples was reverse-transcribed into cDNA using the QuantiTect Reverse Transcription Kit (Qiagen). qPCR was performed using QuantiFast SYBR Green PCR Master Mix (Qiagen) on the ABI 7900HT Sequence Detection System (Life Technologies). Primer sequences were obtained from Primerbank^30^ and purchased from Sigma. Standard curves were prepared from serially diluted RT-qPCR products. Melt-curve analysis was conducted for all samples to confirm the specificity of amplified products. Relative expression levels of target genes were calculated by normalization to the housekeeping genes *HMBS, PPIA* and *PPIB*.^9^ The RT-qPCR was log2 transformed, converted to Z-scores, and analyzed by consensus hierarchical clustering.

### Statistical Analysis

Comparisons of all categorical variables between the clusters were performed using χ^2^ test while ANOVA was used for continuous variables. Kaplan-Meier survival curve was used to assess the time to first re-presentation or admission after recruitment. A p-value of less than 0.05 was considered significant. All analyses were performed using SPSS version 22 (Chicago, Illinois).

## Supplemental Information Titles and Legends

**Figure S1: Identification of IRF7 molecular phenotypes by consensus clustering. A)** The empNegativeControls function from the R package RUVCorr was employed to identify a set of negative controls genes. The plot shows variability on the vertical axis (IQR) versus mean expression on the horizontal axis. The control genes are coloured red. **B)** The RUVnormalise algorithm was employed to model unwanted variation and regress it out of the data. The pairwise correlation between samples was calculated before (left panel) and after (right panel) RUV normalisation. Variations in cellular composition resulted in strong correlation patterns between samples, and this correlation structure was removed by RUV normalisation. **C)** The consensus clustering algorithm calculates a metric called the Cumulative Distribution Function (CDF), which measures consensus across hundreds of different clustering runs for increasing numbers of “k” clusters. The number of clusters in the data is estimated by observing the point at which the CDF reaches a maximum (left plot), and/or as the relative change in area under the CDF curve stabilises (right panel). In this data set, the CDF did not reach a maximum value, but the rate of change of the area under the curve started to plateau at k=5 or k=6. **D)** The consensus cluster analysis result was plotted for k=5 (left panel) and k=6 (right panel). We selected k=5 because the clusters were more well defined.

**Figure S2: Computational inference of the cellular responses underlying IRF7hi and IRF7lo exacerbations.** Relative proportions of 13 major cell types in nasal swab samples collected from children with IRF7hi exacerbations, IRF7lo exacerbations, or from healthy controls. Boxplots showing IQR fenced using the Turkey method. Significant two-way Kruskal-Wallis P values are shown in italics. * represents a Mann-Whitney P value < 0.05, ** < 0.01, *** < 0.001, and **** < 0.0001.

**Figure S3: Computational inference of the cellular responses underlying IRF7hi and IRF7lo exacerbations stratified by RV detection.** Relative proportions of 13 major cell types in nasal swab samples collected from children with IRF7hi exacerbations, IRF7lo exacerbations, or from healthy controls. Boxplots showing IQR fenced using the Turkey method. Significant two-way Kruskal-Wallis P values are shown in italics. * represents a Mann-Whitney P value < 0.05, ** < 0.01, *** < 0.001, and **** < 0.0001.

**Figure S4: Time to next hospital presentation or admission for a respiratory illness comparing children in the discovery cohort with IRF7hi or IRF7lo exacerbations.** Kaplan Meier curve of the proportion of children that have a subsequent hospital presentation/admission for any respiratory diagnosis followed for a maximum of 3.55 years. Censored data represents the length of time a child was followed and did not have another hospital presentation/admission for a respiratory illness.

**Table S1:** Differential gene expression in children with RV positive exacerbations vs RV negative controls.

**Table S2:** Differential gene expression in children with RV positive exacerbations vs RV negative convalescence.

**Table S3:** Differential gene expression in children with RV negative exacerbations vs RV negative controls.

**Table S4:** Differential gene expression in children with RV negative exacerbations vs RV negative convalescence.

**Table S5:** Differential gene expression in children with IRF7hi exacerbations vs RV negative controls.

**Table S6:** Differential gene expression in children with IRF7lo exacerbations vs RV negative controls.

**Table S7:** Molecular drivers of IRF7hi exacerbations.

**Table S8:** Molecular drivers of IRF7lo exacerbations.

**Table S9:** Pubmatrix results.

**Table S10:** Technical variables of samples in the IRF7hi and IRF7lo exacerbations.

